# Protein Language Models Capture Structural and Functional Epistasis in a Zero-Shot Setting

**DOI:** 10.1101/2025.09.14.676130

**Authors:** Ananthan Nambiar, Sayantani B. Littlefield, Carlos Cuellar, Rohit Khorana, Sergei Maslov

**Affiliations:** Department of Bioengineering, University of Illinois Urbana-Champaign; Carl R. Woese Institute for Genomic Biology, University of Illinois Urbana-Champaign; Department of Genome Sciences, University of Washington; eScience Institute, University of Washington; Department of Computer Science, University of Illinois Urbana-Champaign; Center for Biophysics and Quantitative Biology, University of Illinois Urbana-Champaign; Department of Physics, University of Illinois Urbana-Champaign; Theoretical and Computational Biophysics Group, NIH Resource Center for Macromolecular Modeling and Visualization, Beckman Institute for Advanced Science and Technology, University of Illinois Urbana-Champaign

**Keywords:** protein language model, zero-shot, epistasis, protein structure, protein evolution, mutation, variant effect prediction

## Abstract

Protein language models (PLMs) learn from large collections of natural sequences and achieve striking success across prediction tasks, yet it remains unclear what biological principles underlie their representations. We use epistasis, the dependence of a mutation’s effect on its sequence context, as a lens to probe what PLMs capture about proteins. Comparing PLM-derived scores with deep mutational scanning data, we find that epistasis emerges naturally from pretrained models, without supervision on experimental fitness. Raw model scores align with residue–residue contacts, indicating that PLMs internalize structural proximity. Applying a nonlinear transformation to bring model outputs onto the experimental scale, however, shifts the signal toward functional couplings between distant sites. These findings show that PLMs capture both structural and functional dependencies from sequence data alone, and that epistasis provides a powerful window into the biological principles embedded in their representations.

## 1 Introduction

The central idea of Darwinian evolution is that organisms evolve through the accumulation of genetic variations that affect fitness. At the level of proteins, fitness-altering variations often take the form of amino acid substitutions. Some substitutions improve protein stability or function and are positively selected, while others may be deleterious or neutral.

In 1909, British geneticist William Bateson introduced the term epistasis to describe situations where the effect of one genetic change depends on the presence of another [1, 2]. Today, epistasis broadly refers to non-additive interactions between mutations, where the effect of a double mutant is not equal to the sum of the effects of each single mutant (see Figure 1d) [3]. For example, a mutation that is deleterious in one sequence context might be neutral or even beneficial in another. These context-dependent interactions play an important role in shaping protein evolution. They contribute to the ruggedness of protein fitness landscapes, creating local optima and constraining viable evolutionary paths [4, 5]. Understanding and predicting epistasis is therefore a longstanding challenge in molecular biology, with implications for evolutionary theory, protein engineering, and variant interpretation.

**Fig. 1.**
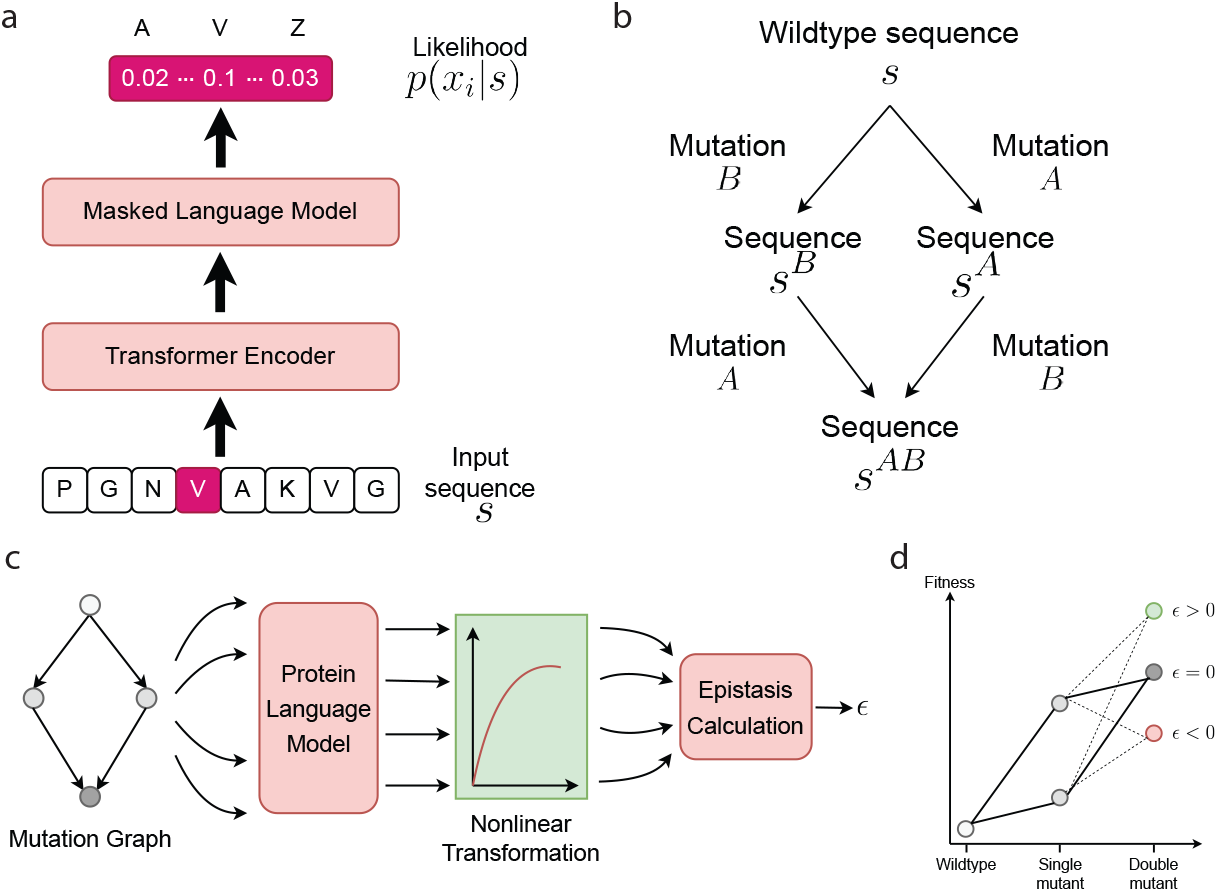
Overview of our framework for assessing epistasis in protein language models. **(a)** A pretrained protein language model (PLM), such as ESM-2, is used to assign relative log-likelihoods to mutations in the context of a given sequence. The model operates on unaligned protein sequences using a masked language modeling objective. **(b)** Mutational space is represented as a directed acyclic graph (DAG) rooted at the wildtype sequence, with branches to single and double mutants. Each double mutant is reachable via two paths, allowing the estimation of conditional mutation effects. These two paths of mutation effects can then be averaged (see Methods) to obtain a single consensus estimate. **(c)** PLM-derived log-likelihoods are transformed via a nonlinear mapping to obtain model-derived fitness values that align with experimental measurements. **(d)** A geometric interpretation of epistasis: if mutation effects are additive, the vector sum of single-mutant fitness values predicts the double-mutant fitness. Deviations from this additive expectation indicate epistasis.

Recent advances in protein language models (PLMs) offer a powerful new approach to describe protein sequences [6, 7]. These models, inspired by their analogs in natural language processing, learn to predict protein sequences by treating them as strings of amino acid tokens.[8, 9]. PLMs are trained in a self-supervised manner to recover masked residues in a sequence based on the surrounding context, allowing them to learn statistical patterns from millions of unannotated protein sequences [8]. This training objective encourages the model to identify which amino acids are most biologically plausible at any particular position in a protein, reflecting evolutionary and biochemical constraints, without access to structural or functional labels.

One practical application of PLMs is variant effect prediction in a zero-shot setting. Because the model assigns likelihoods to amino acid sequences, the relative log-likelihood of a mutant compared to the wildtype can be used as a proxy for its biological fitness [10, 11]. These model-derived scores have been used to design antibodies with improved binding affinity and viral neutralization in just a few rounds of low-throughput screening [12]. These scores have also been used to trace evolutionary trajectories by constructing vector fields over sequence space that recapitulate known evolutionary timelines [13].

However, most such applications assume that the effects of individual mutations are independent and additive. This overlooks the fact that in many proteins, the functional impact of a mutation depends on the surrounding sequence context. In principle, PLMs should be capable of capturing these contextual dependencies, or epistatic interactions, because their training objective is inherently contextual.

In this work, we take a step back and ask: *To what extent do protein language models capture epistasis?* We do not train models to predict epistasis explicitly. Instead, we investigate whether epistatic interactions emerge naturally from the representations learned during pretraining. By comparing PLM-derived fitness scores with experimental measurements of single and double mutants across multiple proteins, we explore how epistasis, and more broadly the interplay between sequence, structure, and function, is encoded in PLMs. Our findings suggest that protein language models encode rich contextual dependencies that reflect both structural and functional sources of epistasis.

## 2 Results

To assess whether protein language models encode epistasis, we devised a framework to compare model-derived fitness scores with experimentally measured fitness across single and double mutants. Our approach proceeds in four steps, illustrated in Figure 1c.

We begin by organizing the mutational space into a directed acyclic graph (DAG) rooted at the wildtype sequence, with branches to mutants (see Figure 1b). In this framework, a given double mutant can arise through two possible paths: mutation *A* followed by mutation *B*, or vice versa. The DAG thus provides a way to compute conditional fitness values for each mutation in the background of the other. We then linearly average the predictions from these two paths to obtain a consistent estimate of the double-mutant fitness. Comparing this estimate to the additive expectation allows us to identify non-additive (epistatic) effects.

Each sequence in the mutation graph is passed through a pretrained PLM, such as ESM-2, to obtain a relative likelihood score, as shown in Figure 1a. Following Rives et al., rather than using the absolute log-likelihood of the mutated sequence, we compute the change in log-likelihood relative to the wildtype sequence [10], defined as

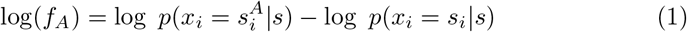

where *s* is the wildtype sequence, *s*_*i*_ is the wildtype residue at position *i* and *s*^*A*^ is the mutant residue at the position where mutation *A* occurs (see Methods for details). This formulation captures the model’s preference for the mutant over the wildtype residue in the given sequence context.

However, we found that the raw relative log-likelihood scores are not directly comparable to experimental fitness values. To address this, we introduce and apply a nonlinear transformation that maps PLM scores to a model-derived fitness scale more closely aligned with experimental effects. Once the transformation is applied, each mutant sequence is associated with a model-derived fitness value, which can be directly compared with experimental measurements or combined to estimate epistatic interactions between mutations.

Finally, we compute pairwise epistasis by comparing the predicted fitness of the double mutant to the additive expectation, from its constituent single mutants (see Methods). If the PLM has internalized residue dependencies during pretraining, these model-derived epistasis estimates are expected to correlate with experimental observations.

In the sections that follow, we quantify the nonlinear relationship between PLM fitness and experimental data (2.1), evaluate how well PLMs recover epistasis (2.2) and investigate how this depends on model size (2.3). Finally, we examine how PLM-inferred epistasis relates to protein structure and function (2.4).

### 2.1 PLM-Derived Fitness Scores Exhibit a Nonlinear Relationship with Experimental Fitness

To ground this analysis, we use three widely studied deep-mutational-scanning datasets—TEM1 *β*-lactamase [14], the YAP1 WW domain [15, 16], and the Pab1 RRM2 domain [17, 18]. Each reports an experimental “fitness,” but the meaning differs: TEM1 reflects growth in antibiotic (a resistance proxy), YAP1 reflects peptide-binding enrichment, and RRM2 reflects rescue of yeast growth when the native gene is repressed. We analyze all datasets on a *log* scale because fitness tends to behave better in log units, and our model outputs are log-likelihoods; when a dataset was already reported in log units, we used it as provided, otherwise we applied a log transform before analysis.

To begin with, we studied how well model-derived fitness scores correspond to experimentally measured fitness values. Mutations can occur either in the wildtype background or in the presence of existing mutations, and we sought to evaluate whether PLM-derived scores reflect fitness in both contexts.

First, for each single mutation *A*, we computed the relative log-likelihood log *f*_*A*_ using a pretrained ESM2-650M model and compared it to the experimental value log 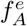 across three deep mutational scanning datasets: TEM1 *β*-lactamase, YAP1, and RRM. As shown in Figure 2a-c, we observe a clear but nonlinear relationship across all three proteins. At low model-derived log-likelihood values predicted by the PLM, the relationship is approximately linear with the logarithm of the experimentally measured fitness, while at higher values this relationship saturates. This pattern highlights the importance of applying a nonlinear transformation to map PLM outputs onto a scale that better reflects biological measurements.

**Fig. 2.**
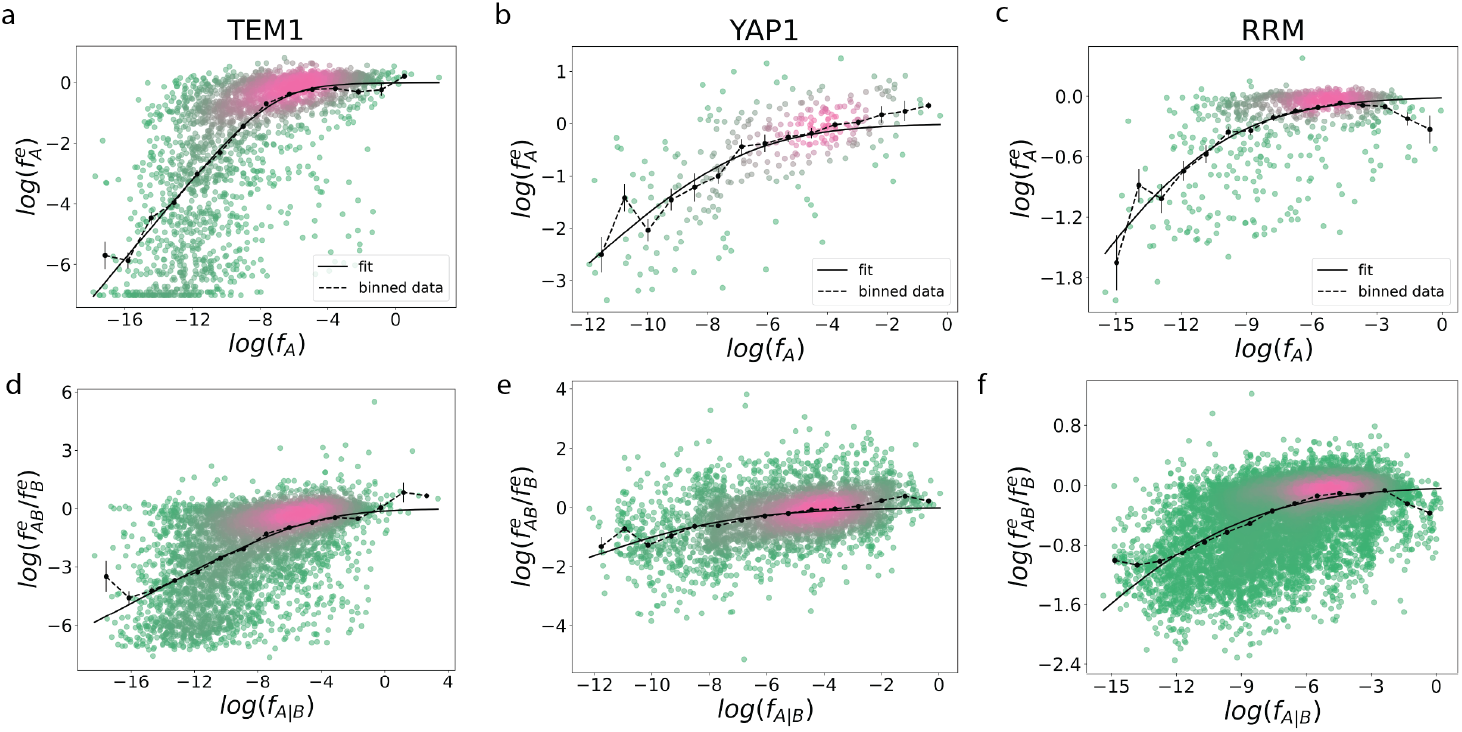
PLM-derived fitness scores show a nonlinear relationship with experimental fitness. **(a–c)** Experimental fitness for single mutations is plotted against PLM-derived relative log-likelihood scores across three proteins: TEM1 *β*-lactamase, YAP1, and RRM. Each point corresponds to a single mutation. Dashed lines show binned averages and solid lines show fits of a nonlinear transformation *ϕ*_1_ used to align model scores with experimental measurements. **(d–f)** Conditional fitness: experimental fitness of mutation *A* in the background of mutation *B*, compared to PLM -inferred log-likelihoods of A given B. Solid lines show fits of a second nonlinear transformation *ϕ*_2_. Both transformations are used to convert PLM scores into modelderived fitness values for epistasis estimation.

To account for this saturation effect, we fit a nonlinear curve of the form

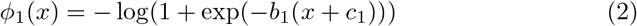

where *b*_1_ and *c*_1_ are fit parameters ((see Methods for details and Supplementary Table 2). This function captures the linear dependence at low values of *x <* −*c*_1_ and the plateau starting for *x > − c*_1_. The solid black lines in Figure 2a–c show the resulting fits, which closely follow the trend of the binned data (dashed lines). We apply this transformation to align PLM scores with experimental fitness in downstream analyses.

We next asked whether PLM-derived scores capture mutation effects not only in the wildtype background, but also in the presence of other mutations. For each pair of mutations *A* and *B*, we computed the fitness of mutation *A* in the background of mutation *B*, denoted log *f*_*A*|*B*_, and compared it to its experimental counterpart. As shown in Figure 2d–f, this conditional comparison reveals a similar but notably different trend from the wildtype background. In particular, the lower tails of the relationship are still linear but less steep, suggesting a narrower dynamic range and reduced sensitivity at the extremes. To account for this, we fit a separate nonlinear transformation, *ϕ*_2_(*x*) with the same functional form as *ϕ*_1_(*x*), but with independently fit parameters *b*_2_ and *c*_2_ (see Methods). The resulting curves (solid black lines) again follow the binned experimental data (dashed lines), allowing us to recover a consistent mapping between PLM scores and experimental fitness even in the presence of background mutations.

We refer to *ϕ*_1_(*x*) and *ϕ*_2_(*x*) as the nonlinear transformation and apply it to all PLM-derived fitness values in downstream analyses.

### 2.2 PLMs Capture Epistatic Interactions After the Nonlinear Transformation

Next, we asked whether protein language models capture epistasis—the interaction between mutations where the effect of one mutation depends on the presence of another. For each pair of mutations *A* and *B*, we define experimental epistasis as

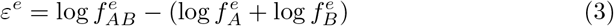

Where 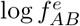 is the experimentally measured fitness of the double mutant and 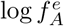 and 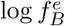 are 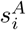 the fitnesses of the corresponding single mutants.

To estimate epistasis from PLMs, we used the mutation graph (Figure 1b) to extract model-derived scores for each mutation, both individually and in the presence of another. For each pair of mutations *A* and *B*, we obtain relative log likelihood scores for *A* in the background of *B*, and vice versa, as well as scores for *A* and *B* in the wildtype background. These scores were then transformed using *ϕ*_1_ and *ϕ*_2_ (defined in Section 2.1) to yield model-derived fitness values. Because a double mutant can be reached through two paths, *A* after *B* or *B* after *A*, we used both to estimate the effect of each mutation in the background of the other. We then compared the average of these conditional fitness values to the sum of their individual effects in the wildtype background, yielding a symmetrized estimate of PLM-inferred epistasis, which we denote by *ε* (See Methods for details). This value reflects the deviation from additivity (i.e., how much the joint effect of two mutations differs from the sum of their individual effects).

As shown in Figure 3a–c, PLM-derived epistasis values are significantly correlated with experimentally measured epistasis across all three proteins, with Pearson *r* = 0.37 for TEM1, *r* = 0.34 for YAP1, and *r* = 0.26 for RRM. PLMs recover a broad spectrum of interaction effects and show consistent alignment with experimental trends despite being evaluated in a zero-shot setting.

**Fig. 3.**
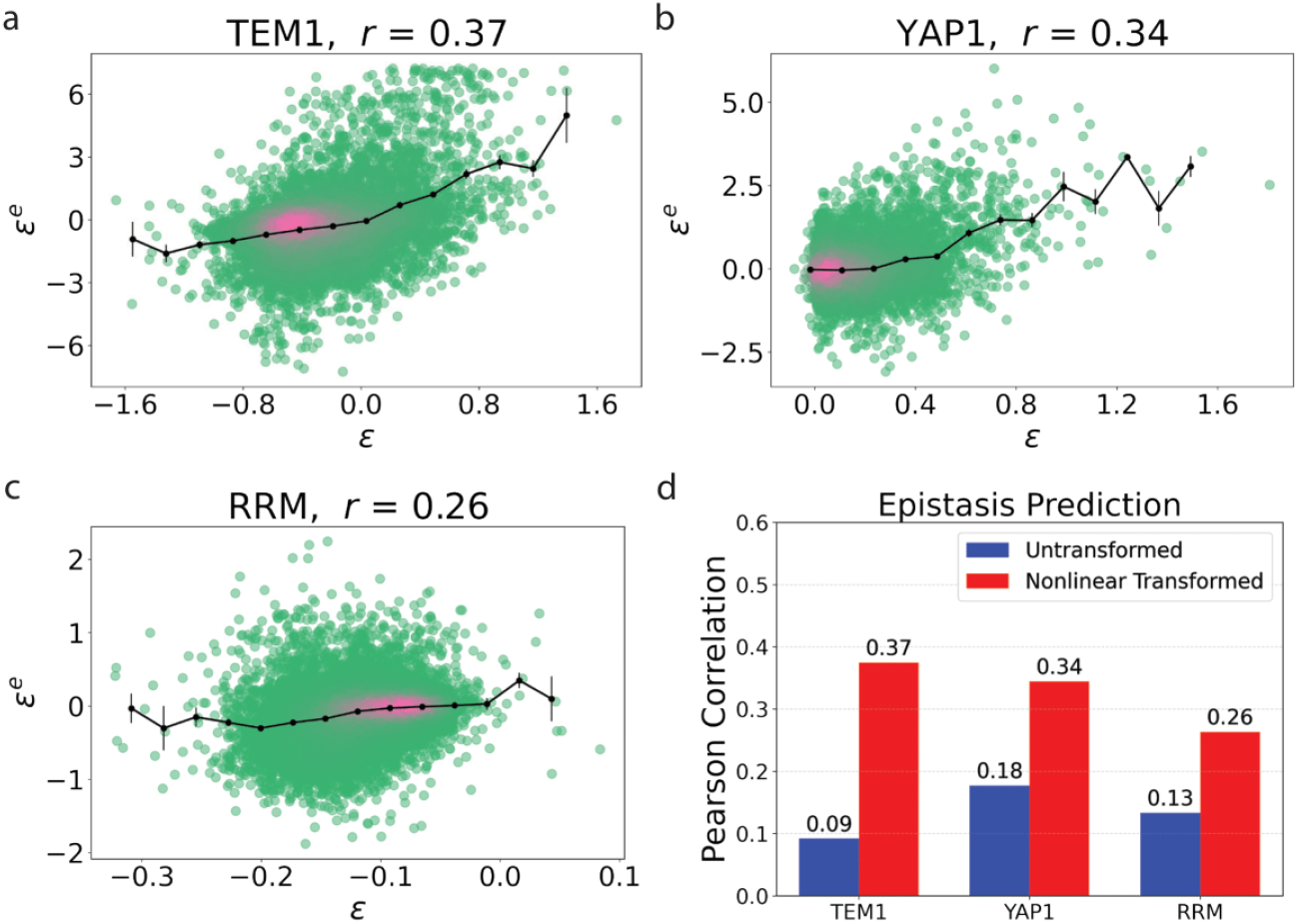
Protein language models recover epistasis in a zero-shot setting. **(a–c)** PLM- derived epistasis estimates *ε* are compared against experimentally measured epistasis *εe*. Each point corresponds to a pair of mutations. Correlations indicate that PLMs recover a wide range of interaction effects without any training on fitness data. **(d)** Effect of the nonlinear transformation: performance drops substantially when epistasis is computed directly from raw log-likelihoods, without applying *ϕ*_1_ and *ϕ*_2_. This highlights the importance of aligning PLM scores with experimental fitness before computing interaction effects.

To assess the impact of the nonlinear transformation, we compared performance with and without applying *ϕ*_1_ and *ϕ*_2_. As shown in Figure 3d, skipping the transformation substantially reduces predictive accuracy, with correlations dropping to *r* = 0.09, 0.18, and 0.13 for TEM1, YAP1, and RRM, respectively. This confirms that the transformation is essential for revealing the epistatic signal embedded in PLM scores.

Together, these results show that PLMs capture not only the individual effects of mutations, but also the dependencies between them. Notably, this epistatic signal emerges naturally from pretraining on large datasets of naturally occurring protein sequences, without any supervision or explicit modeling of interaction terms.

### 2.3 Epistasis and Fitness Prediction Improves with Model Size up to Intermediate Scale

Having established that PLMs can recover epistatic interactions, we then examined how this ability scales with model size. To this end, we evaluated a series of ESM-2 models with parameter counts ranging from 35M to 3B, using the same epistasis prediction pipeline described earlier.

As shown in Figure 4b, epistasis prediction improves with model size up to 650M parameters, where performance peaks across proteins. Pearson correlations reach *r* = 0.38 for TEM1, 0.34 for YAP1, and 0.26 for RRM. Notably, this trend does not hold for the largest model: the 3B-parameter PLM shows a marked drop in performance for TEM1 and YAP1, and no further gain for RRM. This suggests that larger models do not necessarily yield better epistasis predictions and may even overfit to features that are less relevant to interaction effects.

**Fig. 4.**
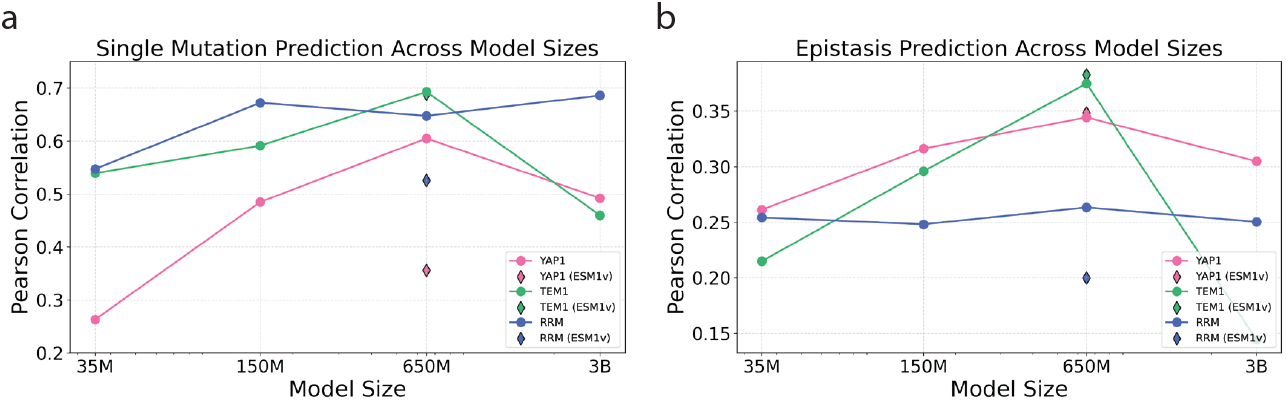
Comparing the predictive abilities of the ESM models model sizes. **(a)** Correlation between PLM-derived and experimental fitness values for single mutations across ESM models of increasing size. **(b)** Correlation between PLM-derived and experimental epistasis across the same model series. Circles indicate ESM-2 models being used while diamonds indicate ESM-1v model predictions.

Figure 4a shows the corresponding results for single-mutation prediction. As with epistasis, performance generally increases with model size up to 650M parameters. Beyond that point, and TEM1 and YAP1 show a performance drop. These results indicate that intermediate-sized PLMs achieve the best performance for zero-shot prediction of epistasis. Beyond this scale, increasing model size does not improve predictions and may even reduce accuracy, potentially due to overfitting to patterns that do not generalize well to mutation-level effects.

We also evaluated the ESM-1v model with 650M parameters, shown as diamonds in Figure 4. For TEM1 and YAP1, ESM-1v slightly outperforms ESM-2 at the same parameter count for epistasis despite ESM-1v underperforming ESM-2 for YAP1.

### 2.4 Nonlinear Calibration Shifts PLM Epistasis from Structural Contacts to Functional Interactions

Building on the nonlinear calibration established in Sections 2.1–2.3, we compare two quantities derived from the same PLM scores: (i) an untransformed epistasis computed directly in log-likelihood space, and (ii) a nonlinearly transformed epistasis computed after mapping PLM scores to model-derived fitness via nonlinear transformations. For each protein, we construct a residue–pair epistasis matrix that summarizes the interaction strength between every pair of positions. To obtain these matrices, we aggregate mutation-level epistasis at the position level. Following [19], we do not distinguish between positive and negative epistasis: for each of the 19 × 19 possible amino-acid substitutions at a residue pair, we sum the squared model-predicted epistasis values and then take the square root to define a single score for that residue pair. Applying this procedure across all residue pairs produces the raw epistasis matrix, *E*_raw_, from untransformed PLM log-likelihoods, and the transformed matrix, *E*, from model-derived fitness values (see Methods).

When *E*_raw_ is compared with the structural contact map, the patterns are strikingly similar: near-diagonal structure and block patterns coincide, indicating that the raw PLM epistasis predominantly reflects geometric/physical proximity and hence structural stability constraints. This matches prior observations that pairwise dependencies inferred from PLMs are enriched for spatial neighbors and contact patterns[19]. After applying the nonlinear transformation, however, the transformed matrix *E* shows a reorganized pattern: banded rows and columns emerge, revealing residues that interact broadly across many partners. These bands identify hubs that often coincide with active-site or binding residues and bridge distant elements, consistent with long-range functional couplings layered on top of structural stability.

Viewed this way, the nonlinear transformation reweights and extends the epistasis signal, shifting emphasis from local, contact-like pairs toward residues that encode biological function in addition to structure. In short, PLM-predicted epistasis computed without the nonlinear map is dominated by structural/stability effects, whereas applying the transformation reveals functional contributions not explained by proximity alone. We now examine these patterns protein-by-protein (TEM1, YAP1, and RRM) showing how untransformed epistasis tracks structural interactions, while transformed epistasis highlights functional couplings between residues.

### 2.4.1 TEM-1 *β*-lactamase

TEM-1 *β*-lactamase enzyme is a prototypical class A *β*-lactamase that confers antibacterial resistance by hydrolyzing the *β*-lactam ring of antibiotics. Its active site is framed by conserved elements with well-established roles: the SDN loop (S130–D131–N132) that helps form the oxyanion hole; the Ω-loop (residues 164 to 179), containing E166, a general base in deacylation; the nucleophilic S70 at the N-terminus of helix H2, crucial for acylation and substrate recognition; and the C77–C123 disulfide bond, which stabilizes helices H3/H4 and maintains the fold [20–23]. We show a structural representation of TEM-1 *β*-lactamase with its most relevant functional regions highlighted (Figure 5d) using VMD to create its cartoon representation [24].

**Fig. 5.**
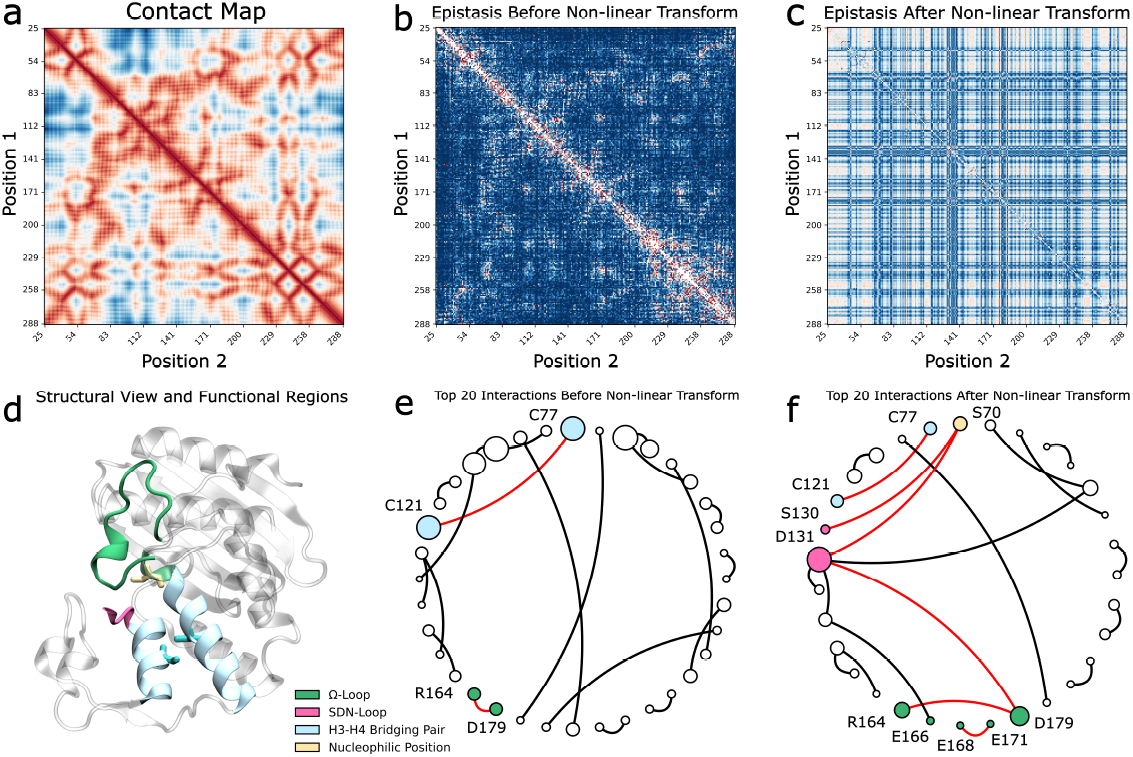
Visualizing the biological underpinnings of PLM-derived epistasis in TEM-1 *β*-lactamase. **(a)** Residue–residue contact map of TEM-1 *β*-lactamase. **(b–c)** Heatmaps of PLM-derived position–pair epistasis matrix (b) before and (c) after applying the nonlinear transformation. **(d)** Cartoon representation of TEM-1 colored by its functional regions. **(e–f)** Before the transformation, epistasis closely mirrors physical contacts in (a). After the transformation, the contact-like signal gives way to a checkerboard-like pattern that highlights distinct epistatic hotspots. Graphs of the top 20 strongest epistatic interactions (by magnitude) before (e) and after (f) transformation. Nodes are residues (scaled by total interaction strength) and edges represent the existence of an epistatic interaction. Node colors indicate key regions: Ω-loop (green), SDN-loop (pink), H3–H4 bridging residue pair (blue), and catalytic S70 (yellow). Only residues that are part of one of the four highlighted regions are labeled. In these circular graphs, residues are arranged around the circumference in sequence order (N→C). Edges that connect functional regions are colored red. Prior to the trainsformation, only the C77–C123 bridge and minor Ω-loop residues (R164, D179) appear among top interactions, whereas after the transformation the network is dominated by functionally critical sites: S70, Ω-loop residues (R164, E166, E168, E171, D179), and the SDN-loop (S130, D131).

Prior to the nonlinear transformation, the position–pair epistasis matrix, *E*_raw_ (Figure 5b) recapitulates the contact map of TEM-1 (Figure 5a), with the strongest interactions confined to pairs of spatially adjacent residues. This correspondence hints that, in its untransformed form, the PLM’s epistatic signals predominantly reflect structural proximity rather than functional coupling. After applying the nonlinear transformation, the epistasis matrix, *E* (Figure 5c) is less aligned with pure contact patterns. Instead, a checkerboard pattern hinting at the existence of epistatic hotspot residues emerges.

Mapping the top 20 epistatic pairs onto interaction graphs clarifies this shift (Figure 5e-f). In these circular graphs, residues are arranged around the circumference in sequence order (N→C), and edges connect residue pairs with high epistasis (node size reflects total interaction strength; see caption). Before the nonlinear transformation (Figure 5e), the only canonical structural or functional motifs highlighted are the C77–C123 disulfide bridge (critical for helix H3/H4 stability[20, 23]) and a faint signature from two Ω-loop residues (R164 and D179), whose interactions remain loop-confined and do not connect to other functional sites in the epistatic network. The majority of strong edges link residues close to each other without known major functional or structural roles[25] and, consistent with the contact-like signal in *E*_raw_, predominantly connect residues that are close in sequence (short arc lengths on the circle) rather than forming long-range links.

Interestingly, after the transformation (Figure 5f) the network reorganizes around experimentally validated functional residues. S70 becomes a prominent epistatic node, reflecting its role in acylation[20, 22]; furthermore, we see it epistatically interact with the SDN-loop and the C77-C123 bridge. The Ω-loop residues R164, E166, E168, E171, and D179 appear prominently and are epistatically linked to the SDN-loop. Simultaneously, SDN-loop positions S130 and D131 (essential for oxyanion stabilization [20, 21]) appear among the top epistatic interactions. After applying the nonlinear transformation, we observe an increase in the amount of structural and functional relevant sites highlighted by epistasis; we also observe functional regions form epistatic connections among them.

### 2.4.2 WW1 domain of YAP1

The WW1 domain of hYAP65/YAP1 is a three–*β*–sheet module that facilitates protein–protein interactions by recognizing proline-rich PPxY motifs through a conserved hydrophobic groove. This groove is formed by a binding region (primarily G188, L190, H192, Q195, T197, and W199) that provides the stacking, hydrogen-bonding, and hydrophobic contacts required for ligand engagement. Within this region, W199 makes an aromatic–proline stacking contact with the first proline of the ligand; T197 and G188 contact the second proline; and L190, H192, and Q195 help position the terminal tyrosine side chain in the pocket [26–28]. At the C-terminus, the functionally important proline P202 supports the fold. Its cyclic structure introduces conformational rigidity that stabilizes the short 3_10_ helix that terminates the WW fold and the local hydrophobic core [28]. Consistent with this role, a P202A substitution has been shown to cause a drastic decrease or loss of binding to PPxY-containing partners such as WBP1 [29]. We show a structural representation of the WW1 domain with its most relevant functional regions highlighted (Figure 6d).

**Fig. 6.**
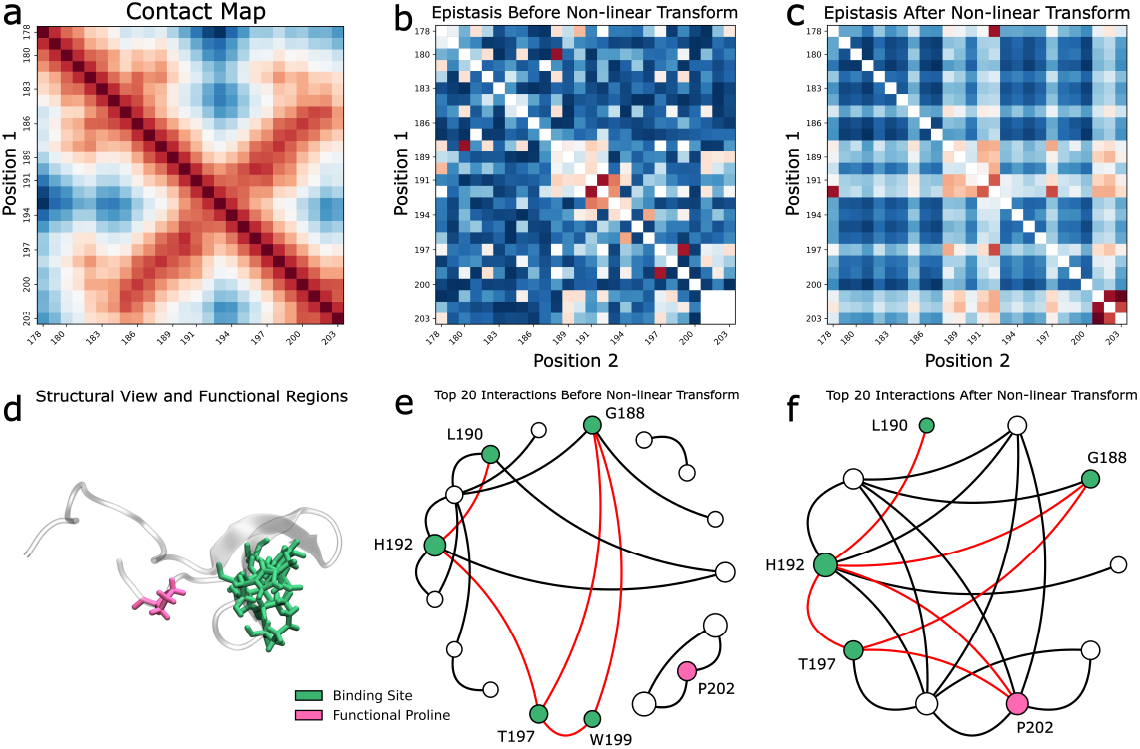
Visualizing the biological underpinnings of PLM-derived epistasis in the WW1 domain of YAP1. **(a)** Residue–residue contact map of the WW1 domain in YAP1. **(b–c)** Heatmaps of PLM-derived position–pair epistasis matrix (b) before and (c) after applying the nonlinear transformation. Before the transformation, epistasis closely mirrors physical contacts in (a). After the transformation, the contact-like signal gives way to a checkerboard-like pattern that highlights distinct epistatic hotspots. **(d)** Cartoon representation of the WW1 of YAP1 domain colored by its functional regions. **(e–f)** Graphs of the top 20 strongest epistatic interactions before (e) and after (f) the transformation. Nodes are residues (scaled by total interaction strength) and edges represent the existence of epistatic interactions. Node colors indicate key regions: WW1 domain’s binding region (green) and functional proline (pink). Only residues that are part of one of the two highlighted regions are labeled.In these circular graphs, nodes are arranged around the circumference in sequence order (N→C). Edges connecting functional residues are colored red. Before the transformation, both regions appear among the top interactions but edges are primarily confined to nearby residues or within each region. After the transformation, the network is dominated by stronger weights on binding-region and P202 nodes, and epistatic links emerge between the two functional regions.

As in TEM-1, the untransformed position–pair epistasis matrix, *E*_raw_ (Fig. 6b), reflects the structure of the contact map (Fig. 6a): the strongest interactions are concentrated near the diagonal and within the binding region, with additional signal between P202 and its immediate neighbors. This correspondence indicates that, prior to the nonlinear transformation, PLM-derived epistasis again largely reflects structural proximity. After applying the transformation, the matrix *E* (Fig. 6c) departs from contact-like structure and exhibits the checkerboard organization, similar to that observed in TEM-1.

Interaction graphs of the top 20 position–pair scores make this shift explicit (Fig. 6e–f). Before the transformation (Fig. 6e), edges are largely confined within regions—within the binding groove and locally around P202. Post-transformation (Fig. 6f), functional hubs dominate and links between them emerge: P202 (magenta) connects to multiple residues in the binding groove, indicating functional coupling between the C-terminal proline and the ligand-recognition pocket.

Together, these results parallel our observations in TEM-1: the untransformed signal is largely contact-like, whereas the transformed epistasis emphasizes functional coupling, connecting P202 with the core binding region of the WW1 domain.

### 2.4.3 RRM2 Motif of PAB1

The second RNA-recognition motif (RRM2) of Pab1 is a canonical ~100-residue RRM fold with a four-stranded antiparallel *β*-sheet packed against two *α*-helices. RNA binding occurs on the exposed *β*-sheet and is dominated by the conserved RNP-1 (on *β*3) and RNP-2 (on *β*1) motifs. Within RNP-1, a group of aromatics (F168, F170, F173) forms a stacking platform for adenines, with F170 and F173 especially sensitive to substitution [18, 30–32] RNP-2 on the other hand, contributes complementary contacts through its consensus positions. We show a structural representation of RRM2 with its most relevant functional regions highlighted (Figure 7d).

**Fig. 7.**
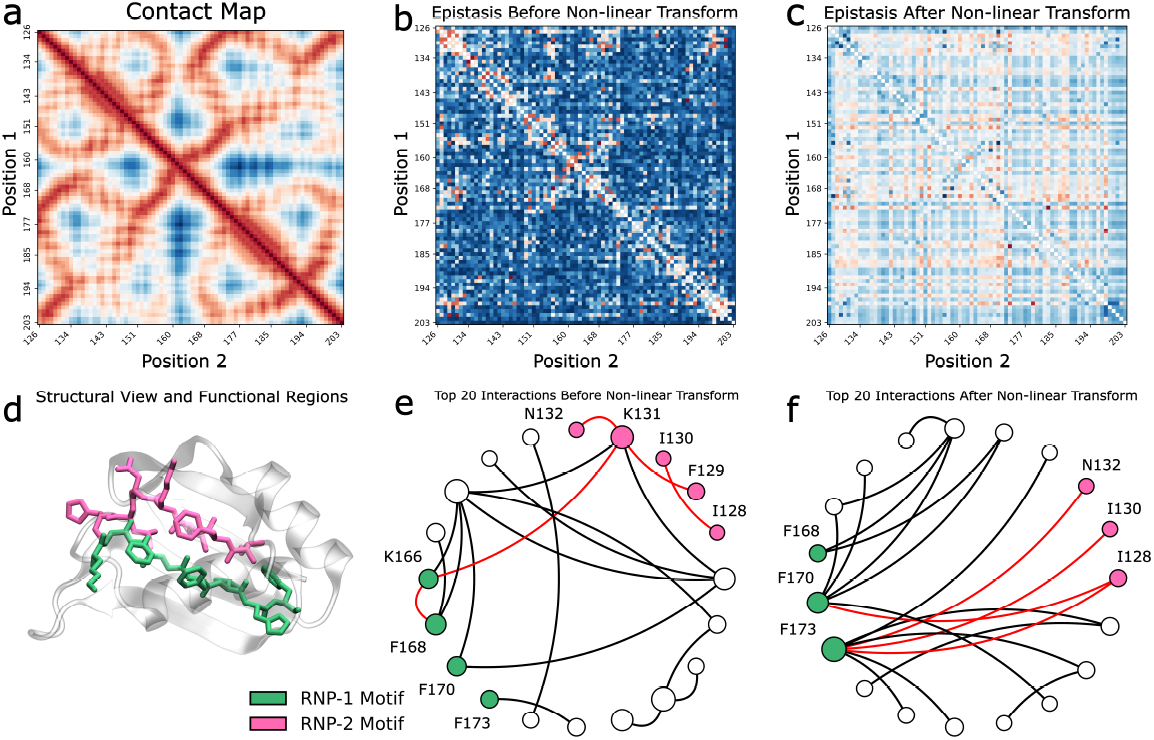
PLM-derived epistasis reflects structural contacts prior to the nonlinear transformation and highlights functional coupling afterward in the RRM2 domain of *S. cerevisiae* Pab1. **(a)** Residue–residue contact map of Pab1 RRM2. **(b–c)** Heatmaps of position–pair epistasis matrix (b) before and (c) after applying the nonlinear transformation. Before the transformation, epistasis closely mirrors physical contacts in (a). After the transformation, the contact-like signal gives way to a checkerboard-like pattern that highlights distinct epistatic hotspots. **(d)** Cartoon representation of RRM2 colored by its functional regions. **(e–f)** Graphs of the top 20 strongest epistatic interactions (by magnitude) before (e) and after (f) NT. Nodes are residues (scaled by total interaction strength) and edges represent the existence of an epistatic interaction. Node colors indicate key regions: RNP-1 motif (green) and RNP-2 motif (magenta). Only residues that are part of one of the two highlighted regions are labeled. In these circular graphs, residues are arranged around the circumference in sequence order (N→C). Edges involving at least one residue from a highlighted region are colored blue. Prior to the transformation, both motifs appear among the top interactions but edges are largely confined within each motif and the aromatic F residues of RNP-1 are not emphasized. After the transformation, the network is dominated by RNP-1 aromatic nodes (F168, F170, F173), and epistatic links emerge between these RNP-1 residues and the RNP-2 motif.

As in the other proteins, the untransformed position–pair matrix *E*_raw_ (Fig. 7b) looks similar to the the contact map (Fig. 7a), although it does struggle to capture the structure which arises from the stacked *β*-sheet formation of the RRM2 motif. However, it is still consistent with a primarily proximity-driven readout as observed in the other proteins before the nonlinear transformation. After applying the nonlinear transformation, the matrix *E* (Fig. 7c) once again displays the checkerboard pattern like with TEM-1 and the WW domain of YAP1.

The interaction graphs tell a similar story (Fig. 7e–f). Without the transformation, edges are mostly confined to nearby residues. Post-NT, the network re-centers on the RNP-1 aromatics: F170 and F173 emerge as prominent hubs (with F168 also present), and cross-motif links from RNP-1 to RNP-2 become prevalent. Thus, the transformed signal emphasizes the RNP-1 stacking platform and its coupling to RNP-2, aligning with residues directly responsible for poly(A) recognition. While the contrast between the untransformed and transformed graphs are the least stark in RRM2, compared to TEM-1 and YAP1, this is because the key functional motifs of RRM2 (RNP-1 and RNP-2) are within the domain’s *β*-sheet structure, which provides structural rigidity to the domain. Moreover, ≈ 83% of mutations in RRM2 are deleterious [33], making it difficult to distinguish functionally essential sites from regions important primarily for structural stability.

Together with the TEM-1 and YAP1 results, this pattern reinforces that the nonlinear transformation shifts the PLM signal from contact-like proximity to functionally organized coupling between key binding motifs. Structural views support this interpretation (Figure S1a–c). In TEM-1 *β*-lactamase (Figure S1a), the molecular surface colored by each residue’s maximum transformed epistasis magnitude shows hotspots that coincide with the SDN loop (S130–D131–N132), the Ω-loop (164–179), the catalytic S70, and the C77–C123 disulfide bridging H3–H4. This co-localization confirms that the transformed signal prioritizes functionally consequential positions, shifting from a contact-like pattern to a network emphasizing functional hubs and cross-motif coupling. Structural visualization supports this interpretation for the WW1 domain of YAP1 as well (Figure S1b): the transformed epistasis hotspots align with known functional and structurally relevant motifs and are concentrated in the binding groove and at P202. Finally, the structural view for Pab1 RRM2 (Figure S1c) again supports the interpretation that the nonlinear transformation highlights functional regions.

## 3 Discussion

In this study, we investigate whether pretrained PLMs contain information about epistasis, how the effect of a mutation depends on its sequence context, and how to turn that information into biologically meaningful quantities. Unlike most zero-shot variant-effect work with PLMs that evaluates only single substitutions or assumes additivity, we ask if interaction effects emerge from sequence alone and whether a simple calibration can reveal the non-additive couplings that biologists care about. Concretely, we compute epistasis from PLM log-likelihoods and introduce a monotone nonlinear transformation that aligns these scores to experimental fitness while preserving rank order.

Across three unrelated proteins, we find that epistasis is indeed present in PLMs without any supervision on fitness: pairwise interaction terms computed directly from raw log-likelihoods correlate with experimentally measured epistasis. Importantly, applying a simple monotone nonlinear transformation to the PLM scores further improves agreement with experiment. This establishes that context dependence is encoded in pretrained models and can be exposed with light calibration.

To understand what the calculated epistasis represents biologically, we first compared the position–pair signal to structural contact maps. Using the untransformed epistasis calculation, the position–pair signal visually mirrors contact maps, indicating that models internalize local geometric constraints. This is consistent with observations from Zhang et al., who showed that PLM-derived pairwise dependencies recover spatial neighbors via a “categorical Jacobian”, which is an analog to our matrix *E*_raw_ [19]. After calibration, the organization changes: signals concentrate around functional hubs and inter-motif links, such as catalytic-loop couplings in TEM-1 (Figure 5) and connections between the C-terminal proline and the WW binding groove in YAP1 (Figure 6). The nonlinear transformation step is, to our knowledge, new, and across the three proteins, this shift from structure to function is reproducible and coincides with improved agreement with experimental epistasis.

We also explored how model scale affect epistasis recovery. Performance improves with model size up to an intermediate scale (on the order of hundreds of millions of parameters) and then plateaus or slightly declines. One possibility is that very large models overfit distributional quirks in training corpora that are orthogonal to mutation-level fitness effects. Practically, “bigger” is not always “better” for epistasis; medium-scale models strike a favorable balance of accuracy and cost.

These findings could have future applications. First, prospective library design could use epistasis maps to prioritize double mutants with predicted compensation or synergy, shrinking combinatorial libraries. Moreover, mechanistic mapping could highlight long-range couplings and functional clusters using calibrated epistasis. This, in turn could help researchers better understand how a protein functions by focusing experiments on epistatic hubs identified by PLMs. To enable these applications in practice, a minimal workflow is to (1) experimentally measure a small calibration panel of single-mutation effects and a focused set of conditional effects (A in the background of B); (2) fit the nonlinear mappings (*ϕ*_1_) and (*ϕ*_2_) on those measurements; (3) predict PLM-derived fitness and epistasis for candidate pairs; and (4) use the predicted epistasis to rank pairs depending on the desired application. While our study did not set out to optimize such workflows, future work may refine and benchmark them for practical use.

Moreover, this study also opens up opportunities for further studies in understanding how PLMs capture epistasis. For example, we evaluate only pairwise interactions. However, higher-order effects likely contain additional structure that is currently not captured by our approach. Future work could also move beyond pairs by extending our directed-acyclic-graph framework to triples and higher-order sets to infer epistatic hypergraphs.

This work also raises questions about why a nonlinear transformation is needed and why a separate nonlinear transformation is required for mutations in the background of a first mutation. Empirically, the raw PLM scores are linearly correlated with experimental fitness for deleterious mutations (negative fitness), but beneficial or compensatory mutations with positive fitness collapse into a plateau (Fig. 2a–c). This could because PLM likelihoods reflect not only biophysical fitness but also evolutionary accessibility and historical sampling. Variants easier to reach—for example, amino acid changes accessible by a single nucleotide step—are overrepresented in natural sequence corpora and thus assigned higher likelihoods, even when their assay fitness is comparable to less accessible alternatives. If so, this bias would flatten the upper tail of the distribution and necessitate a nonlinear calibration *ϕ*_1_ to restore sensitivity to positive-fitness mutations. The need for a second nonlinear transformation in mutated backgrounds stems from the fact that double mutants create sequence contexts rarely observed in natural corpora. In mutated backgrounds, the sequence context can be out-of-distribution for the PLM. Double mutants create residue combinations and long-range states that are rare in nature, introducing a need for a separate transformation. In practice, the conditional calibration *ϕ*_2_ has a shallower slope in its quasi-linear regime (Fig. 2d–f), reflecting the compressed dynamic range of PLM likelihoods under out-of-distribution conditions. Future work could delve into understanding how the pretraining corpus affects how the PLM captures epistasis. This could take the form of disentangling the contributions of evolutionary accessibility and biophysical fitness, or investigating how PLMs behave with out-of-distribution sequences.

Taken together, our results show that pretrained PLMs encode both structural and functional dependencies from sequence alone. A simple, biologically motivated nonlinear calibration exposes long-range couplings that align with catalytic and binding motifs, enabling practical gains for design and interpretation. More broadly, epistasis exposes the biological content of PLMs, illuminating how sequence alone captures structure and function.

## 4 Methods

### 4.1 Obtaining Relative Log-Likelihoods from PLMs

Let *s* = (*s*_1_, …, *s*_*L*_) denote the wild-type (WT) amino acid sequence. For a single mutation *A* that changes position *i* from *s*_*i*_ to 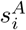, we compute the relative loglikelihood (RLL) log *f*_*A*_ as defined in Eq. (1), i.e., using the difference of the model’s position-wise log-probabilities for the mutant versus the WT residue at position *i* in the WT background. For a second mutation *B* that changes position *j* from *s*_*j*_ to 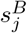, we compute the conditional score log *f*_*A*|*B*_ by evaluating Eq. (1) in the background where *B* is present, *s*^*B*^ instead of the wild-type sequence.

### 4.2 Nonlinear Transformation of Relative Log-Likelihoods

We convert relative log-likelihood (RLL) scores from the PLM into model-derived fitness values using the monotone nonlinearity specified in Eq. (2), where we fit the parameters (*b, c*) by nonlinear least squares.

For each protein, we define the calibration subset by a 20% uniform random sample of the double-mutant entries. The remaining 80% of double mutants are held out and used only for downstream evaluation of PLM-derived epistasis and all figures/tables that report generalization. To fit the single-mutant curve, we take the set of single substitutions that appear as constituents of the double mutants in the 20% calibration subset.

#### Single-mutant curve

We fit one instance of Eq. (2) to unique single mutants by regressing experimental single-mutant fitness (on the log scale) against PLM RLLs for the corresponding substitutions using a 20% fitting subset. This yields parameters (*b*_1_, *c*_1_) and a mapping *ϕ*_1_ that transforms single-mutant RLLs log *f*_*A*_ into model-derived fitness values. The same (*b*_1_, *c*_1_) are used for all positions and amino-acid identities (i.e., a global curve for single mutants).

#### Conditional (background) curve

Separately, we fit a second instance of Eq. (2) to *conditional* targets derived from double-mutant measurements using the same 20% fitting subset. For each pair (*A, B*), we form the background-adjusted log-fitness contrasts

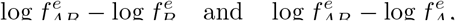

and regress them on the corresponding conditional PLM scores log *f*_*A*|*B*_ and log *f*_*B*|*A*_, respectively. This produces parameters (*b*_2_, *c*_2_) and a mapping *ϕ*_2_ that transforms conditional RLLs into background-specific model-derived fitness terms. As with the single-mutant curve, (*b*_2_, *c*_2_) are shared globally across all positions and mutation identities.

### 4.3 Epistasis Calculation

With the single-mutant and conditional transformations *ϕ*_1_ and *ϕ*_2_ defined above, we quantify epistasis between mutations *A* and *B* by averaging the two mutation-order paths on the model-derived fitness scale and subtracting the additive singlemutant baseline:

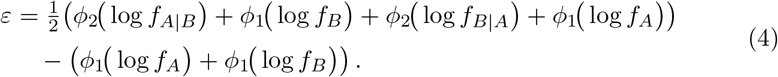

Essentially, the two path totals (see Figure 1b) *P*_*A*→*B*_ = *ϕ*_1_(log *f*_*A*_) + *ϕ*_2_(log *f*_*B*|*A*_) and *P*_*B*→*A*_ = *ϕ*_1_(log *f*_*B*_) + *ϕ*_2_(log *f*_*A*|*B*_) are averaged symmetrically, and the additive expectation *ϕ*_1_(log *f*_*A*_) + *ϕ*_1_(log *f*_*B*_) is subtracted. By construction, *ϵ* is symmetric in *A* and *B*.

### 4.4 Calculating the position–pair score

We go over all combinations of residue pairs (*i, j*). For each such pair, we enumerate all possible amino-acid substitutions at *i* and *j* (i.e., *A* ∈ ℳ_*i*_, *B* ∈ ℳ _*j*_), evaluate the epistasis *ε*(*A, B*) for that ordered pair using the symmetric two-path construction defined above, and collect the resulting values across all (*A, B*).

We then collapse these mutation-level values to a single position–pair score via a root–sum–of–squares (RSS) aggregation:

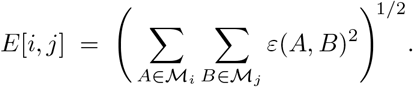

On the pre-transformation (PLM) scale, we repeat the exact procedure—computing epistasis using the raw scores prior to applying Eq. (2) to obtain *E*raw.

### 4.5 Data Sources

Across all three datasets, we retained only single- and double-substitution measurements and recomputed experimental pairwise epistasis from the reported single- and double-mutant fitness values using the definition shown on Equation 3. To place measurements on a comparable scale, if a dataset reported fitness on a log scale, we used those values as provided; otherwise, we applied a log transform to the reported fitness prior to analysis.

#### TEM-1 β-lactamase

We used the mutation dataset from Gonzalez and Ostermeier [14]. This study reports fitness values for 12k sequential (consecutive-position) double mutants and 5,460 single mutants, with fitness normalized to the wild type. In line with our uniform processing across proteins, we retained only single and double amino-acid substitutions with associated experimental fitness measurements and recomputed experimental pairwise epistasis.

#### YAP1 WW domain

We used the processed deep mutational scanning dataset released by Chen *et al*. [15], which aggregates measurements from Araya *et al*. [16]. This dataset contained a total of 8,670 pairs of mutations which were from various positions in the protein. In the original paper, the fitness score quantifies binding of WW domain variants to a cognate peptide ligand at high throughput (deep-sequencing-based enrichment).

#### Pab1 RRM2

For the second RNA-recognition motif (RRM2) of the *S. cerevisiae* poly(A)-binding protein (Pab1), we used the processed dataset compiled by Rollins *et al*. [17], derived from the deep mutational scan of Melamed *et al*. [18]. This dataset includes 36,500 pairs of amino acid mutations and the corresponding experimentally measured fitness for single and double mutations. The original experiment scored variants by their ability to support yeast growth in a PAB1-complementation assay. For the wild-type sequence, we used the sequence provided by UniProt [34].

## Code and Data

The code and data to reproduce these results are available at https://github.com/maslov-group/Epistasis

## Acknowledgements

This work utilizes resources supported by the National Science Foundation’s Major Research Instrumentation program, grant #1725729, as well as the University of Illinois at Urbana-Champaign for the usage of HAL [35]. Part of this work was performed under the auspices of the U.S. Department of Energy by Argonne National Laboratory under Contract DE-AC02-06-CH11357. A.N. would like to thank members of the Noble Lab for useful discussions.

## Author Contributions

A.N. and S.M. devised and supervised the study; A.N., S.B.L., C.C. and R.K. performed data analysis; all authors discussed and wrote the paper.

## Declarations

The authors declare no competing interests.

## 5 Supplementary

**Table S1.** Pearson correlation coefficients between experimental and predicted fitness for single mutations

| Protein | Untransformed | Transformed |
| --- | --- | --- |
| TEM-1 (Beta lactamase) | 0.6501 | 0.6903 |
| YAP1 (WW1) | 0.6578 | 0.6668 |
| RRM (Pab1) | 0.5228 | 0.6060 |

**Table S2.** Nonlinear fit parameters for and correlation coefficients between experimental and computational epistasis

| TEM-1 | Untransformed | $b_2 = b_1, c_2 = c_1$ | $(b_1, c_1)$ and $(b_2, c_2)$ fit separately |
| --- | --- | --- | --- |
| $b_1$ | - | 0.6639 | 0.6639 |
| $c_1$ | - | 7.2371 | 7.2371 |
| $b_2$ | - | 0.6639 | 0.4298 |
| $c_2$ | - | 7.2371 | 4.6874 |
| Pearson Coefficients | 0.0918 | 0.1169 | 0.3748 |
| YAP1 domain | Untransformed | $b_2 = b_1, c_2 = c_1$ | $(b_1, c_1)$ and $(b_2, c_2)$ fit separately |
| $b_1$ | - | 0.5492 | 0.5492 |
| $c_1$ | - | 7.2231 | 7.2232 |
| $b_2$ | - | 0.5492 | 0.4354 |
| $c_2$ | - | 7.2231 | 8.7439 |
| Pearson Coefficients | 0.1770 | 0.2364 | 0.3444 |
| RRM domain | Untransformed | $b_2 = b_1, c_2 = c_1$ | $(b_1, c_1)$ and $(b_2, c_2)$ fit separately |
| $b_1$ | - | 0.3512 | 0.3512 |
| $c_1$ | - | 11.7190 | 11.719 |
| $b_2$ | - | 0.3512 | 0.3027 |
| $c_2$ | - | 11.7190 | 10.4994 |
| Pearson Coefficients | 0.1333 | 0.1508 | 0.2633 |

**Fig. S1.**
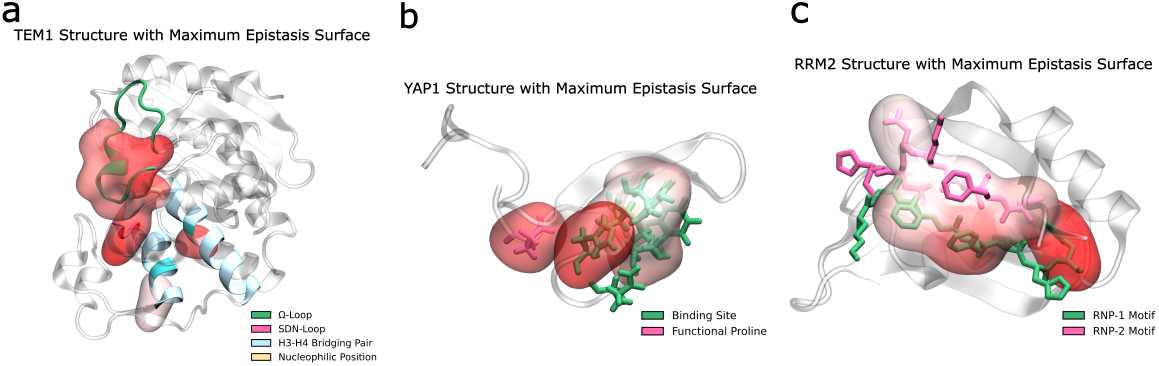
Surface views colored by per-residue maximum transformed epistasis highlight functional hotspots. For each protein, the molecular surface is restricted to residues belonging to functional regions that exhibit strong epistatic coupling in the transformed epistasis graphs (Figs. 5f, 6f, 7f), and is colored by the per-residue maximum transformed epistasis value. A full cartoon of the respective structures is shown inside each surface for structural context, with the highlighted regions indicated. **(a)** TEM-1 *β*-lactamase: hotspots map to the SDN loop, the Ω-loop, the catalytic S70, and the H3–H4 disulfide bridge. **(b)** WW1 domain of YAP1: hotspots concentrate in the ligand-binding groove and at P202. **(c)** Pab1 RRM2: hotspots localize to the RNP-1/RNP-2 motifs.

## Notes

### Competing Interest Statement

The authors have declared no competing interest.

## References

[1] Phillips, P.C.: Epistasis—the essential role of gene interactions in the structure and evolution of genetic systems. Nature Reviews Genetics 9(11), 855–867 (2008)

[2] Bateson, W.: Mendel’s Principles of Heredity. Cambridge University Press, Cambridge [Eng.] (1913)

[3] Lipsh-Sokolik, R., Fleishman, S.J.: Addressing epistasis in the design of protein function. Proceedings of the National Academy of Sciences 121(34), 2314999121 (2024)

[4] Starr, T.N., Thornton, J.W.: Epistasis in protein evolution. Protein science 25(7), 1204–1218 (2016)

[5] De Visser, J.A.G., Krug, J.: Empirical fitness landscapes and the predictability of evolution. Nature Reviews Genetics 15(7), 480–490 (2014)

[6] Koo, P.K., Dallago, C., Nambiar, A., Yang, K.K.: Machine Learning for Protein Science and Engineering. Cold Spring Harbor Lab (2025)

[7] Heinzinger, M., Rost, B.: Artificial intelligence learns protein prediction. Cold Spring Harbor Perspectives in Biology 16(9), 041458 (2024)

[8] Rives, A., Meier, J., Sercu, T., Goyal, S., Lin, Z., Liu, J., Guo, D., Ott, M., Zitnick, C.L., Ma, J., et al.: Biological structure and function emerge from scaling unsupervised learning to 250 million protein sequences. Proceedings of the National Academy of Sciences 118(15), 2016239118 (2021)

[9] Nambiar, A., Heflin, M., Liu, S., Maslov, S., Hopkins, M., Ritz, A.: Transforming the language of life: transformer neural networks for protein prediction tasks. In: Proceedings of the 11th ACM International Conference on Bioinformatics, Computational Biology and Health Informatics, pp. 1–8 (2020)

[10] Meier, J., Rao, R., Verkuil, R., Liu, J., Sercu, T., Rives, A.: Language models enable zero-shot prediction of the effects of mutations on protein function. Advances in neural information processing systems 34, 29287–29303 (2021)

[11] Brandes, N., Goldman, G., Wang, C.H., Ye, C.J., Ntranos, V.: Genome-wide prediction of disease variant effects with a deep protein language model. Nature Genetics 55(9), 1512–1522 (2023)

[12] Hie, B.L., Shanker, V.R., Xu, D., Bruun, T.U., Weidenbacher, P.A., Tang, S., Wu, W., Pak, J.E., Kim, P.S.: Efficient evolution of human antibodies from general protein language models. Nature biotechnology 42(2), 275–283 (2024)

[13] Hie, B.L., Yang, K.K., Kim, P.S.: Evolutionary velocity with protein language models predicts evolutionary dynamics of diverse proteins. Cell systems 13(4), 274–285 (2022)

[14] Gonzalez, C.E., Ostermeier, M.: Pervasive pairwise intragenic epistasis among sequential mutations in TEM-1 β-lactamase. Journal of molecular biology 431(10), 1981–1992 (2019)

[15] Chen, L., Zhang, Z., Li, Z., Li, R., Huo, R., Chen, L., Wang, D., Luo, X., Chen, K., Liao, C., et al.: Learning protein fitness landscapes with deep mutational scanning data from multiple sources. Cell Systems 14(8), 706–721 (2023)

[16] Araya, C.L., Fowler, D.M., Chen, W., Muniez, I., Kelly, J.W., Fields, S.: A fundamental protein property, thermodynamic stability, revealed solely from large-scale measurements of protein function. Proceedings of the National Academy of Sciences 109(42), 16858–16863 (2012)

[17] Rollins, N.J., Brock, K.P., Poelwijk, F.J., Stiffler, M.A., Gauthier, N.P., Sander, C., Marks, D.S.: Inferring protein 3D structure from deep mutation scans. Nature genetics 51(7), 1170–1176 (2019)

[18] Melamed, D., Young, D.L., Gamble, C.E., Miller, C.R., Fields, S.: Deep mutational scanning of an RRM domain of the Saccharomyces cerevisiae poly(a)-binding protein. RNA 19(11), 1537–1551 (2013)

[19] Zhang, Z., Wayment-Steele, H.K., Brixi, G., Wang, H., Kern, D., Ovchinnikov, S.: Protein language models learn evolutionary statistics of interacting sequence motifs. Proceedings of the National Academy of Sciences 121(45), 2406285121 (2024)

[20] Jelsch, C., Mourey, L., Masson, J.-M., Samama, J.-P.: Crystal structure of Escherichia coli TEM1 β-lactamase at 1.8 °A resolution. Proteins: Structure, Function, and Bioinformatics 16(4), 364–383 (1993)

[21] Minasov, G., Wang, X., Shoichet, B.K.: An ultrahigh resolution structure of TEM-1 β-lactamase suggests a role for Glu166 as the general base in acylation. Journal of the American Chemical Society 124(19), 5333–5340 (2002)

[22] Egorov, A., Rubtsova, M., Grigorenko, V., Uporov, I., Veselovsky, A.: The role of the Ω-loop in regulation of the catalytic activity of TEM-type βlactamases. Biomolecules 9(12) (2019)

[23] Pimenta, A.C., Fernandes, R., Moreira, I.S.: Evolution of drug resistance: Insight on TEM β-lactamases structure and activity and β-lactam antibiotics. Mini-Reviews in Medicinal Chemistry 14(2), 111–122 (2014)

[24] Humphrey, W., Dalke, A., Schulten, K.: VMD: Visual molecular dynamics. Journal of Molecular Graphics 14(1), 33–38 (1996)

[25] Majiduddin, F.K., Materon, I.C., Palzkill, T.G.: Molecular analysis of beta-lactamase structure and function. International Journal of Medical Microbiology 292(2), 127–137 (2002)

[26] Sudol, M., Shields, D.C., Farooq, A.: Structures of YAP protein domains reveal promising targets for development of new cancer drugs. Seminars in Cell Developmental Biology 23(7), 827–833 (2012)

[27] McDonald, C.B., McIntosh, S.K.N., Mikles, D.C., Bhat, V., Deegan, B.J., Seldeen, K.L., Saeed, A.M., Buffa, L., Sudol, M., Nawaz, Z., Farooq, A.: Biophysical analysis of binding of WW domains of the YAP2 transcriptional regulator to PPXY motifs within WBP1 and WBP2 adaptors. Biochemistry 50(44), 9616–9627 (2011)

[28] Martinez-Rodriguez, S., Bacarizo, J., Luque, I., Camara-Artigas, A.: Crystal structure of the first WW domain of human YAP2 isoform. Journal of

[29] Espanel, X., Sudol, M.: Yes-associated protein and p53-binding protein2 interact through their WW and SH3 domains*. Journal of Biological Chemistry 276(17), 14514–14523 (2001)

[30] Brambilla, M., Martani, F., Bertacchi, S., Vitangeli, I., Branduardi, P.: The Saccharomyces cerevisiae poly (A) binding protein (Pab1): Master regulator of mRNA metabolism and cell physiology. Yeast 36(1), 23–34 (2019)

[31] Cusick, M.E.: RNP1, a new ribonucleoprotein gene of the yeast Saccharomyces cerevisiae. Nucleic Acids Research 22(5), 869–877 (1994)

[32] Deardorff, J.A., Sachs, A.B.: Differential effects of aromatic and charged residue substitutions in the RNA binding domains of the yeast poly(a)-binding protein. Journal of Molecular Biology 269(1), 67–81 (1997)

[33] Young, D.L.: High throughput determination of sequence-function relationships in protein and RNA. Ph.D. dissertation, University of Washington (2016)

[34] The UniProt Consortium: PABP YEAST (P04147), Polyadenylate-binding protein. https://www.uniprot.org/uniprotkb/P04147/entry. Accessed: 2025-07-17 (2025)

[35] Kindratenko, V., Mu, D., Zhan, Y., Maloney, J., Hashemi, S.H., Rabe, B., Xu, K., Campbell, R., Peng, J., Gropp, W.: HAL: Computer system for scalable deep learning. In: Practice and Experience in Advanced Research Computing, pp. 41–48 (2020)

